# Host-pathogen Immune Feedbacks Can Explain Widely Divergent Outcomes from Similar Infections

**DOI:** 10.1101/2021.01.08.425954

**Authors:** Stephen P. Ellner, Nicolas Buchon, Tobias Dörr, Brian P. Lazzaro

## Abstract

A longstanding question in infection biology is why two very similar individuals, with very similar pathogen exposures, may have very different outcomes. Recent experiments have found that even isogenic *Drosophila melanogaster* hosts, given identical inoculations of some bacterial pathogens at suitable doses, can experience very similar initial bacteria proliferation but then diverge to either a lethal infection or a sustained chronic infection with much lower pathogen load. We hypothesized that divergent infection outcomes are a natural result of mutual negative feedbacks between pathogens and the host immune response. Here we test this hypothesis *in silico* by constructing process-based dynamic models for bacterial population growth, host immune induction, and the feedbacks between them, based on common mechanisms of immune system response. Mathematical analysis of a minimal conceptual model confirms our qualitative hypothesis that mutual negative feedbacks can magnify small differences among hosts into life-or-death differences in outcome. However, explaining observed features of chronic infections requires an extension of the model to include induced pathogen modifications that shield themselves from host immune responses at the cost of reduced proliferation rate. Our analysis thus generates new, testable predictions about the mechanisms underlying bimodal infection outcomes.

## 1 Introduction

Despite more than a century of infectious disease research, we still do not understand why two similar individuals exposed to nearly identical bacterial infections may experience dramatically different outcomes, with some dying while others mount a successful defense and survive. It is routine to define the LD_50_ of a given pathogen as the infectious dose at which half the infected hosts will die. But why do half die while the other half survive? Analogously, we have very little understanding of why some individuals develop severe infections while others remain safe and healthy after similar exposures to opportunistic pathogens. Widely divergent outcomes, even when controlling for genotype and environment, give the appearance that the outcome is random or arbitrary.

We have recently found that *Drosophila melanogaster* reared in a common controlled environment experience biphasic outcomes after identical injections (insofar as experimentally possible) of an opportunistic pathogen [7]. Some hosts die from from acute infection with a high pathogen burden, others survive infection but sustaining a lifelong chronic bacterial burden at much lower density (Fig. 1A). That pattern occurs even when the hosts are isogenic (Fig. 1B). Very similar patterns are seen in *Drosophila* infected with other bacteria [6, 14], flour beetles (*Tribolium castaneum*) infected with *Bacillus thuringiensis*, with higher survival among offspring of immune-primed mothers (Fig. 1C, data from [20, Fig.1D]), virus-infected flies [8], and *Plasmodium*-infected mosquitos (Fig. 1D, data from [1, Fig. 2A]).

**Figure 1:**
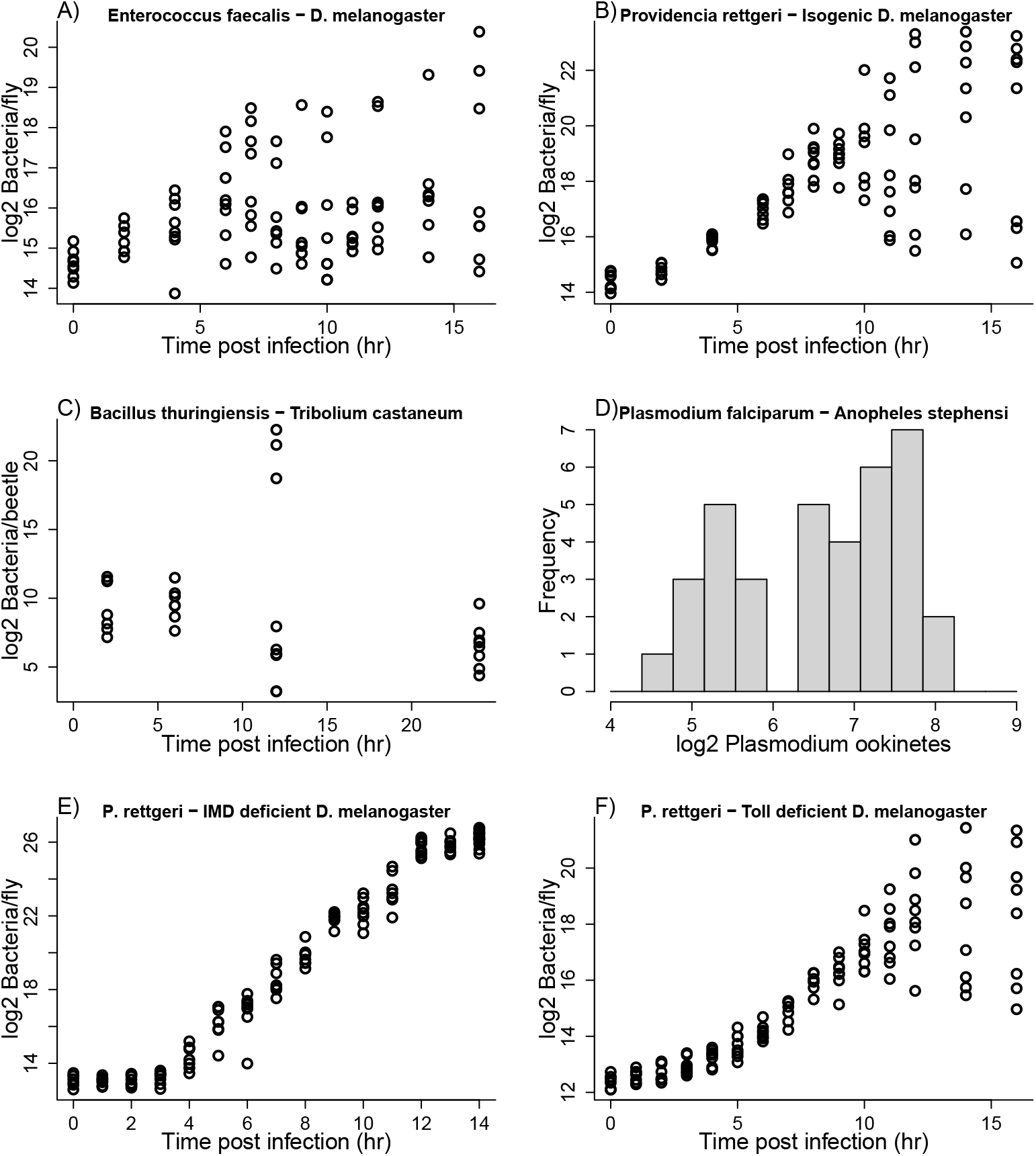
**A),B),E),F)** Some experimental results from Duneau et al. [7] in which flies *D. melanogaster* were given uniform injections of opportunistic pathogens but infection outcomes could be highly variable. Each data point represents a fly that was sacrificed some time after infection to assay its total bacterial load. **C)** Experimental results from [20] in which flour beetles *Tribolium castaneum* were experimentally infected with *Bacillus thuringensis*. The data plotted are unprimed beetles from Experiment 1 of that paper. The high bacterial load beetles at 12h post infection were described as “moribund” [20]. **D)** Experimental results from Bian et al. [1, Fig. 2A] in which mosquitos *Anopheles stephensi* were experimentally infected with *Plasmodium falciparum*. The plotted data are *Plasmodium* ookinetes per midgut lumen in the LBT mosquito strain. Figure drawn by script Figure 1.R.

**Figure 2:**
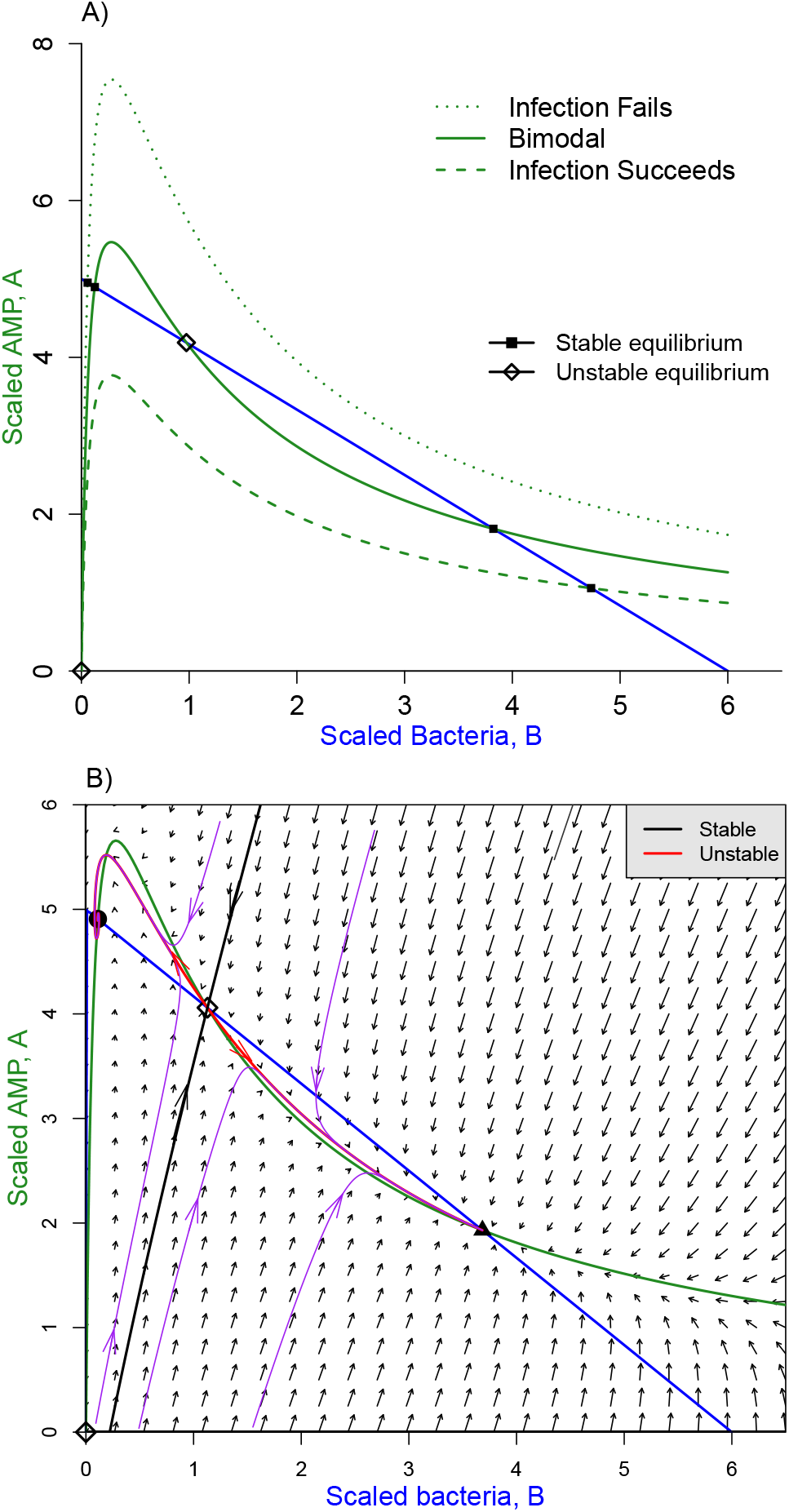
Phase-plane diagrams of the conceptual model. **A)** Possible nullcline configurations. The blue line is the *B* nullcline, the three green curves are the *A* nullcline for three different values of *Q_A_* (4, 5.8, and 8) from lowest to highest, with *K* = 6, *δ_A_* = 0.05, *c* = 0.2, *m* = 0.45 Equilibria (where nullclines intersect) can be stable (solid square) or unstable (open diamond). **B)** Phase portrait in the bistable case. Black and red curves are the stable and unstable manifolds of the interior unstable equilibrium, which is a saddle. Solution trajectories (purple curves) converge to one or the other stable equilibia, depending on the location of their starting point lies. Figures were created by script files `BAnullclinesPlot.R, BAModel.R`.

Production of anti-microbial peptides (AMP) is a principal defense against invading bacteria in *Drosophila* and many other insects [13]. AMP production following pathogen invasion may be up-regulated primarily through the Imd or Toll signaling pathways (or both in combination), depending on the structure of the peptidoglycan in the bacterial cell wall [3]. Response to *Providencia rettgeri* primarily involves the Imd pathway. Flies deficient in Imd-dependent immune response all experienced lethally high pathogen burdens following inoculation with *P. rettgeri* (Fig. 1E), while bimodal infection outcomes persisted in Toll-deficient mutants (Fig. 1F) and in phagocytosis-deficient mutants [7].

In attempting to explain the observed bimodal outcomes, Duneau et al. [7] therefore tested whether flies vary in the speed and magnitude of Imd pathway induction. They found substantial variation in mRNA levels of the *Diptericin* gene, a readout of Imd pathway activity, 4 hours after pathogen injection. At that time, which is prior to the divergence in outcomes, Imd activity was more variable than bacterial load. Thus, the Imd variability presumably reflects intrinsic variability among the flies (despite their genetic homogeneity and common rearing), rather than being a side-effect of differences in bacterial population growth. Based on that finding, Duneau et al. [7] presented a phenomenological model positing that a fly either succeeds or completely fails to control the infection, depending on whether Imd up-regulation occurs before or after bacterial density crosses some threshold. Bimodality of outcomes is thus an *assumption* of their model, not an outcome.

Our goal here is to develop a general mechanistic explanation for outcome bimodality as an emergent property of interactions between the pathogen and host immune responses. van Leeuwen et al. [21] have recently presented an explanation specifically for intestinal parasites, based on nutritional interactions between parasite and host [e.g., 9, 10]. The mechanism in their model, parameterized for a nematode parasite of mouse, is competition for energy and nutrients: a larger pathogen population is increasingly able to divert the resources ingested by the host from the host to itself. The pathogen thus benefits from increased abundance (an Allee effect), potentially resulting in bimodal outcomes where infection duration is long or short depending on whether the initial pathogen abundance is above or below a threshold.

Here we propose an alternative, broadly applicable explanation, that bimodal infection outcomes are a natural result of two negative feedbacks: hosts mount an immune response to eradicate the pathogen, while pathogens attempt to counteract or squelch the immune responses so they can proliferate at the expense of the host. Then, depending on the balance between host immune response and pathogen counter-response, the outcome can be bistable dynamics, in which similar initial states lead to widely divergent outcomes. A simple analogy is the well-known “toggle switch” model for two genes that mutually repress each other’s activity levels. For suitable parameter values, this results in two stable equilibria (each with one gene “on” and the other “off”), separated by an unstable saddle equilibrium. Two trajectories with very similar initial conditions near the origin, but on opposite sides of a separatrix (the stable manifold of the saddle) follow similar paths initially but then separate and eventually converge to different stable equilibria. Continuous variation in initial conditions spanning the separatrix produces discrete variation in outcomes.

We first present a minimal conceptual model for our hypothesis based on the *Drosophila* experimental system. We posit that flies respond to a bacterial infection by producing bacteriocidal AMPs, while bacteria can inactivate AMPs by sequestration and produce proteases that degrade AMPs [12]. In addition, bacteria can produce effectors that interfere with AMP production [e.g., 16]. A slow-fast approximation to this model produces a two-dimensional system, and phase plane analysis of that system verifies our hypothesis that bimodality is a robust outcome of the mutual negative feedbacks.

Importantly, we do not merely confirm that the “toggle switch” mechanism for bistability can be made to operate in a host-pathogen interaction. Our analysis shows that bistability occurs in our model across a wide range of biologically plausible parameter values, and it identifies several specific scenarios in which small between-host differences can be amplified into widely divergent outcomes.

However, analysis of the minimal model shows that for biologically reasonable parameter values, the “toggle switch” mechanism does not provide a complete explanation for the experimental observations. Specifically, it cannot explain the common observation that the pathogen is controlled in surviving hosts but not eliminated or reduced to very low numbers. Rather, there is a a chronic infection held in check by sustained immune system activation [5], which can break out into an active infection if the host immune response is subsequently eliminated (B.P. Lazzaro, *unpublished data*). We therefore extend the model by allowing bacteria to enter a “protected” state where they are partially shielded from immune response. Several such mechanisms for bacterial defense against AMPs are known, including biofilm formation and various cell envelope modifications [12]. The conditions for stable chronic infection in the extended model lead to new, testable predictions about the mechanisms that account for chronic infection rather than complete or near-complete elimination of the pathogen.

Finally, we develop a detailed model for the IMD signaling pathway, to show that our minimal model’s “cartoon” description of immune system activation dynamics, and how it varies among individuals, can be realized in a completely mechanistic model for a defense activation pathway.

## 2 Conceptual model

Our conceptual model tracks a bacterial population *B* growing within an invertebrate host, suffering mortality caused by host-produced AMPs *A*, and producing proteases *R* that degrade AMPs:

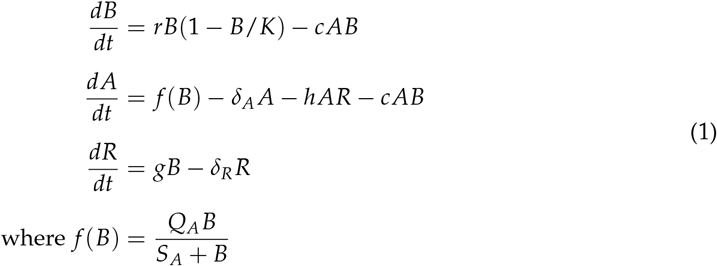

and all parameters are positive. In the absence of AMPs, bacteria have logistic population growth with maximum per-capita growth rate *r* and “carrying capacity” *K*. The carrying capacity corresponds to pathogen growth ceasing because the host is completely consumed, so any model solution where *B* gets close to *K* is interpreted as pathogen killing the host. *cAB* is bacteria mortality due to AMPs. AMP production rate *f* is a function of bacteria abundance, monotonic increasing from *f*(0) = 0 and saturating at maximum rate *Q_A_. S_A_* is the bacterial abundance at which AMP production rate reaches half its maximum value. In our model, AMPs are lost three ways: natural degradation at rate *δ_A_A*, degradation by protease at rate *hAR*, and sequestration, i.e. each “kill” of a bacterium binds and thus inactivates the *A* molecule that was involved. Over the time-scale of interest AMPs are very stable molecules, so *δ_A_* ≪ 1 [17,18]. However, AMPs can be produced quickly enough to create a lethal within-host environment for the pathogen [3]. Proteases *R* are produced by bacteria at constant per-capita rate *g* and degrade naturally at rate *δ_R_R*, which is not necessarily very small. Protease is not consumed in the process of promoting AMP degradation, so the *hAR* term is not replicated in the *dR/dt* equation.

To keep this “proof of concept” model as simple as possible, we have omitted two potential features of the host-pathogen interaction: a constitutive immune response (i.e., production of AMPs in the absence of bacteria [11]), and bacterial production of effectors that interfere with the host mounting an immune response [e.g., 16] rather than acting through AMP degradation and sequestration. In Electronic Supporting Material ESM S.2 we present an extended model that includes these features. We show that the only qualitative effect of the extensions is to add one more scenario (described at the end of this section) where small individual differences in the host immune induction can produce bimodal outcomes.

Before analysis we re-scale the model, setting 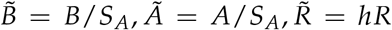 and 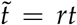. For the sake of visualization and analysis, we reduce the dimension of the model by assuming that *R* is a “fast” (i.e., *g* and *δ_R_* are large) that remains close to its steady state conditional on the other variables, *R* = *mB* where *m* = *g*/*δ_R_*. The calculations are carried out in Maxima script `RescaleBAR.max`. Then dropping the tilde’s on rescaled variables and parameters for clarity, the model we consider is

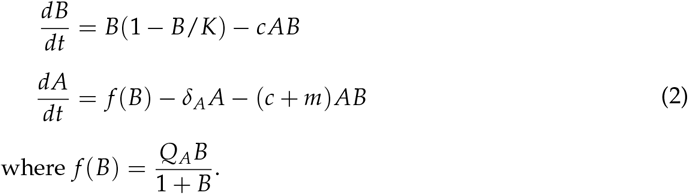

Note that equilibria 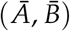 of the reduced model (2) are in 1-to-1 correspondence with equilibria 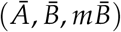 of the full system (1). Because bacterial abundance is now scaled relative to the half-saturation abundance for immune response, and immune response is triggered when bacteria are far below a lethal abundance, we can assume that *K* ≫ 1. Time is scaled so that bacteria that are unhindered by resource shortage or immune response would double in log 2 ≈ 0.7 time units. Observed doubling times are typically on the order of 1h in real time [7], so we can still assume that *δ_A_* is a small parameter in the rescaled model.

Equilibria occur at intersections of the *B* and *A* nullclines (the sets of (*B, A*) values at which *dB/dt* = 0 and *dA/dt* = 0, respectively). The *B* nullcline consists of the axis *B* = 0 and the line *A* = *c*^−1^ (1 – *B/K*); the *A* nullcline is the curve

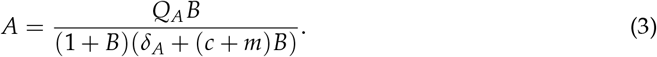

The infection-absent state (*B* = *A* = 0, open diamond) is always an equilibrium. Analysis of the model (in electronic supplementary material ESM S.1) shows that this equilibrium is always unstable: a small inoculum of bacteria initially increases. Other equilibria and their stability depend on the configuration of the *B* and *A* nullclines in the interior of the first quadrant. There are three possibilities, shown in Figure 2 A). Model behavior in each of these cases is analyzed in electronic supplementary material ESM S.1. When host immune response is very strong (large *Q_A_*, dotted green curve) there is only one nullcline intersection giving a stable equilibrium at *B* ≈ 0 and *A* ≈ 1/*c*. Model solutions starting anywhere except *B* = *A* = 0 converge to that equilibrium: immune response always holds the infection in check. When host immune response is very weak (small *Q_A_*, dashed green curve) there is again only one possible outcome: the stable equilibrium near *B* = *K*, *A* = 0, representing a pathogen that has overcome the host’s immune defenses. In between these extremes (solid green curve) there are three interior equilibria, one unstable and two locally stable, at widely differing pathogen densities.

Figure 2 B) illustrates how, in the three equilibrium case, small differences in initial conditions can produce large differences in the infection outcome. The unstable manifold of the middle interior equilibrium consists of two solution trajectories with exactly the right initial conditions so that solutions converge to the middle equilibrium. Initial conditions off the stable manifold lead to infection dynamics that first approach the middle equilibrium, but then veer off to one of the stable interior equilibria, depending on which side of the stable manifold they started. The right-most equilibrium is always a node, but the left-most can be a spiral (as in this example) if it occurs where the *A* nullcline is very steep.

In electronic supplementary material ESM S.1 we derive the following approximate condition for occurence of bistability in the scaled model:

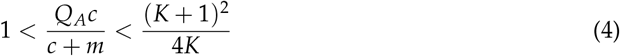

Although eqn. (4) is approximate, we have found numerically that bistability occurs when neither inequality is close to being violated. This condition can be interpreted biologically, showing that the requirements for bistability will often be satisfied. In the scaled model, *K* is the pathogen carrying capacity relative to the half-saturation constant for immune system up-regulation. Thus, *K* is large, and the right-most term will be large, so long as the host responds strongly to a bacterial infection when it is still far below the level at which the host’s survival is threatened. The middle term can be written as *Q_A_* / (1 + *m/c*). *Q_A_* determines how quickly the host can produce AMPs to ward off a pathogen attack, and *m*/*c* is a measure of how effectively the pathogen can counter the host by degrading AMPs, relative to the lethality of the host response. The middle term is thus a measure of the “balance of forces” between host and pathogen – if either antagonist is too strong or too weak, there is only one possible outcome. Condition (4) thus says that if the pathogen is a sufficient threat that the host responds to its presence in low numbers, bistability will occur across a wide range of values for the “balance of forces” between host and pathogen.

The location of the stable manifold depends on parameter values. Here, parameter values were such that small differences in initial bacterial density produce radically different outcomes. In the Duneau et al. [7] experiments, where flies differed in the time required for immune activation, the “initial bacterial density” would be the bacterial density at the time of immune activation, with higher bacterial density after a longer delay. The dynamics in Fig. 2 B) thus provide a qualitative explanation for observed bimodality in outcomes from very similar inoculations of very similar flies.

Figure 3 shows simulations of experiments like those in Fig. 1 using the minimal model. We assumed that host individuals varied randomly in their pattern of immune system activation. This variability could have several different biological causes, including host resource or energetic limitations, but at the level of our model all that matters is the temporal pattern of activation. In all hosts, following the pathogen inoculation, the AMP production rate term *f*(*B*) was multiplied by a three-piece function representing immune system activation: a delay period during which AMP production rate is zero; a linear ramp-up from zero to one; and thereafter constant at 1. The duration (in hours) of the delay period was chosen from a Uniform[1.5, 2.5] distribution, and the time required for the ramp-up was chosen from a Uniform[1,2] distribution. Each plotted point represents one simulated host that was “sacrificed” at a random time, and Gaussian “measurement error” with *σ* = 0.35 was added to log_2_(*B*). These simulations show that our conceptual model provides a potential mechanistic basis for the hypothesis of Duneau et al. [7] that small variations in the speed of immune system activation can produce drastic, bimodal variation in outcomes. In Electronic Supplementary Material ESM S.3 we develop a detailed mechanistic model for the Imd signaling pathway leading to AMP production, and confirm that among-individual variation in kinetic parameters of the pathway can produce a wide range of temporal patterns for immune activation (Fig. S-4).

**Figure 3:**
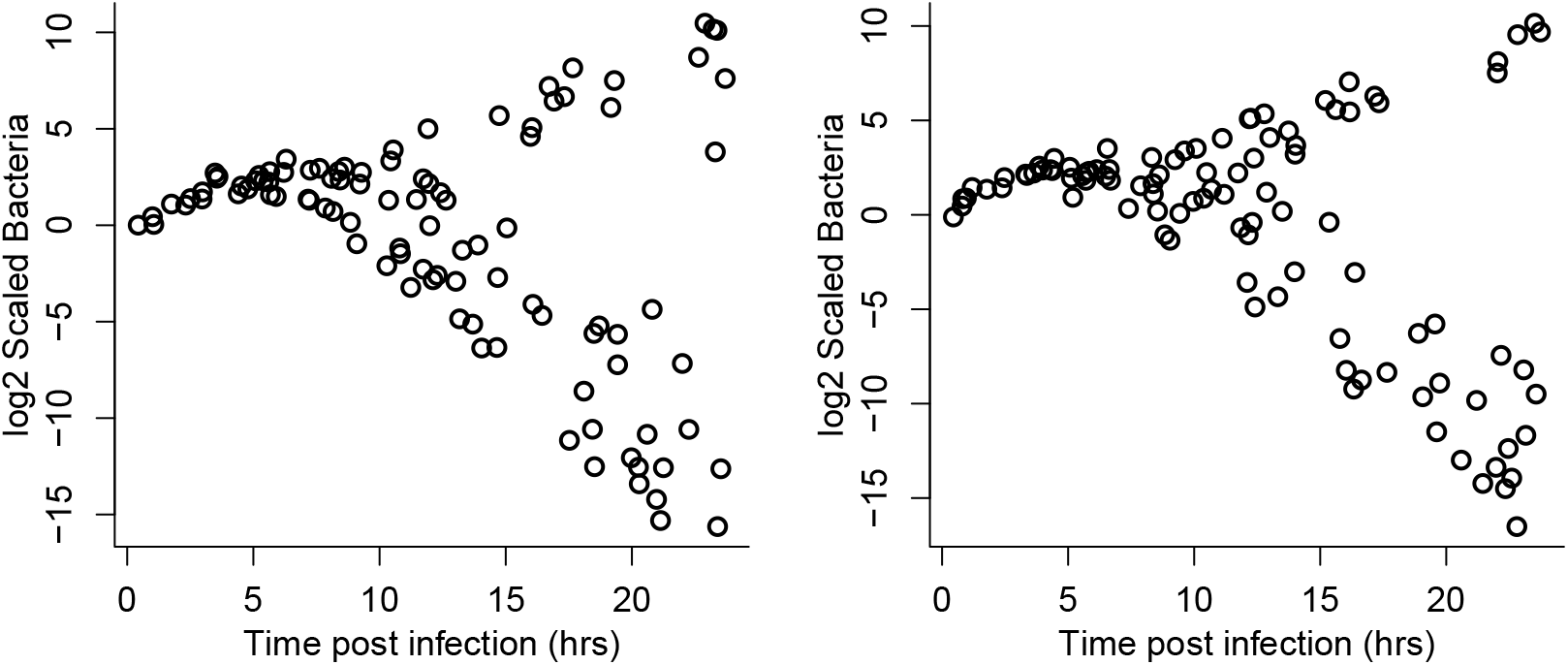
Two replicates of a simulated experiment with 100 host individuals each in the minimal model eqn. (2), with parameter values *Q_A_* = 10, *δ_A_* = 0.02, *c* = 0.1, *m* = 0.2, *K* = 1000. For comparison with experimental results, time was not scaled; bacteria had intrinsic growth rate *r* = 0.5 multiplying the logistic growth term. Each plotted point represents “data” on one host individual. Hosts vary in the delay period and ramp-up speed to full immune response, as described in the main text. Hosts were “sacrificed” at a random time between 0.25 and 24 hours, and Gaussian “measurement error” with *σ* = 0.35 was added to log_2_(*B*). Figure made by script `Split_Outcomes_BA.R.`

At other parameter values, the lower left branch of the unstable manifold approaches the *A* axis, rather than the *B* axis. In our extended model that includes constitutive defense (see Electronic Supplementary Material ESM S.2) this situation creates another way for small between-host differences to produce bimodal outcomes. Constitutive defense moves the pathogen-free equilibrium from the origin to a point (0, *α*) on the *A* axis. The location of this equilibrium relative to the stable manifold determines whether a small invading pathogen population sparks a lethal infection, or is driven down by the host immune response (fig S-2B,C). This scenario can produce bimodal outcomes from small variance among hosts in their constitutive defense levels.

## 3 Chronic infection and protected pathogens

The minimal model can explain bimodal outcomes, but cannot explain another important experimental observation: that the alternative to host death may be a chronic, low-level infection where bacteria remain present at substantial but non-lethal levels, and the host immune response is never fully down-regulated [7, 5]. At the low-B equilibrium in fig. 2 the pathogen density is extremely low. This is not just a feature of the particular parameters in that figure. The slope of the *A* nullcline (green curve) at *B* = 0 is *Q_A_/δ_A_*, so it rises very steeply, and the peak of the nullcline occurs at 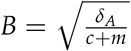. So under the biologically relevant assumption that *δ_A_* is small, and host and pathogen interact strongly (so *c* and *m* are not small), the low-*B* equilibrium will always occur at very low *B*. At the parameters used in Fig. 3, the low-*B* equilibrium is very near zero even though *δ_A_* is not greatly smaller than *c* or *m*. In our extended model (Electronic Supplementary Matrial ESM S.2) the low-*B* equilibrium can be at *B* = 0 when hosts have constitutive AMP production. However, empirical observation is that substantial bacterial loads can persist for the duration of life in hosts that survive the initial infection [7], engendering only a mild reduction in lifespan [4]. Moreover, as bacterial abundance has been scaled in the model so that *S_A_* = 1, *B* ≪ 1 implies that the immune system is almost completely down-regulated, which is also out of line with the experimental observations.

To remove this conflict with empirical observations, we add one more feature to the pathogen population model: the ability of pathogens to achieve some degree of protection from the host immune response, at the cost of reducing their division rate. Several mechanisms are known that can produce this effect [12]. One is for cells to enter a “tolerant” or “persister” state, analogous to known mechanisms of antibiotic tolerance [2] involving either physiological changes (such as cell wall reduction or loss) or a reduction in metabolic rate (dormancy, Westblade et al. [22]). A second is for pathogens to invade some tissue that is protected from the host immune response. For example, intracellular pathogens such as *Mycobacterium tuberculosis* (the causative agent of tuberculosis) and *Listeria monocytogenes* do this by allowing themselves to be phagocytosed, then living inside the macrophage while being protected from host immune responses [15]. Any tissue isolated from the host immune system could play the same function. A final possibility is for cells to form a structure such as a biofilm that protects most cells against host immune response, and allows them to safely remain metabolically active to some extent, while not changing dramatically in numbers [12]. For our purposes we need not distinguish between these possible mechanisms; we can just posit that cells can activate some mechanism affording protection at the cost of reduced proliferation rate.

We thus extend the model to distinguish between “normal” bacteria *N*, and “protected” bacteria *P*. For a minimally complex proof of concept model, we assume that protected cells are completely invulnerable to AMPs (without specifying how this is achieved), but have a lower intrinsic division rate and protease production rate than normal bacterial (by a factor *η* < 1), and a lower carrying capacity *L* (lower carrying capacity is a necessary assumption in this model, as slower division would otherwise simply delay the progression to a high lethal pathogen burden, but host defenses could also limit protected bacterial growth if the assumption of complete invulnerability is relaxed). We assume that the per-capita conversion rate from *N* to *P* states is a sigmoid increasing function *p*(*A*) of AMP concentration. Because our focus is on modeling chronic infection states where the immune system remains activated, we omit back-conversion from *P* to *N* states that might occur at low AMP concentrations. However, we do assume that division of *P* cells produces both a fraction *ν* of *N* cells. When AMP concentrations are low, these *N* cells could proliferate and potentially re-seed a growing infection.

The model is then

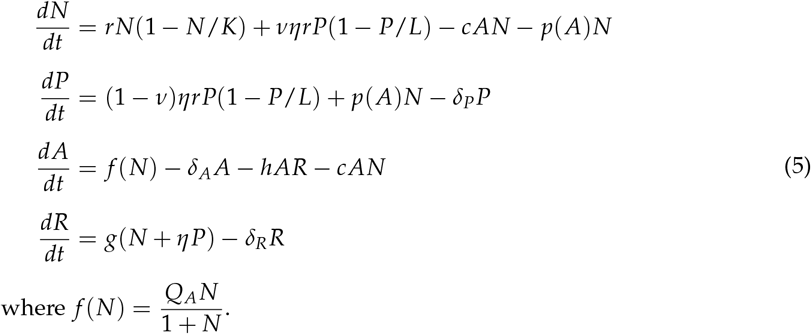

For this section we again scale state variables so that *S_A_* = *h* = 1, but leave time unscaled for the sake of comparisons with experimental results. A doubling time of 1 - 2 hours (*r* = 0.35 – 0.7) can be taken as typical for the *Drosophila* pathogens shown to exhibit bimodal infection outcomes [7].

The extended model readily produces bimodal outcomes in which the pathogen is never reduced to extremely low levels (Fig. 4). Simulations where the pathogen is held in check (fig. 5) confirm that the model can capture the known qualitative features of chronic infections: the outcome is a stalemate, converging to a stable equilibrium where a small bacterial population that continues to undergo cell divisions is held down by host immune responses. For these parameters there are enough *N*-state bacteria, sustained by division of *P*-state cells, to keep AMP production at roughly 30% of its maximum rate. However, sustained AMP production could also result, in theory, if protected bacteria do not divide but nonetheless produce metabolic products that induce a host immune response (having little or no effect on them).

**Figure 4:**
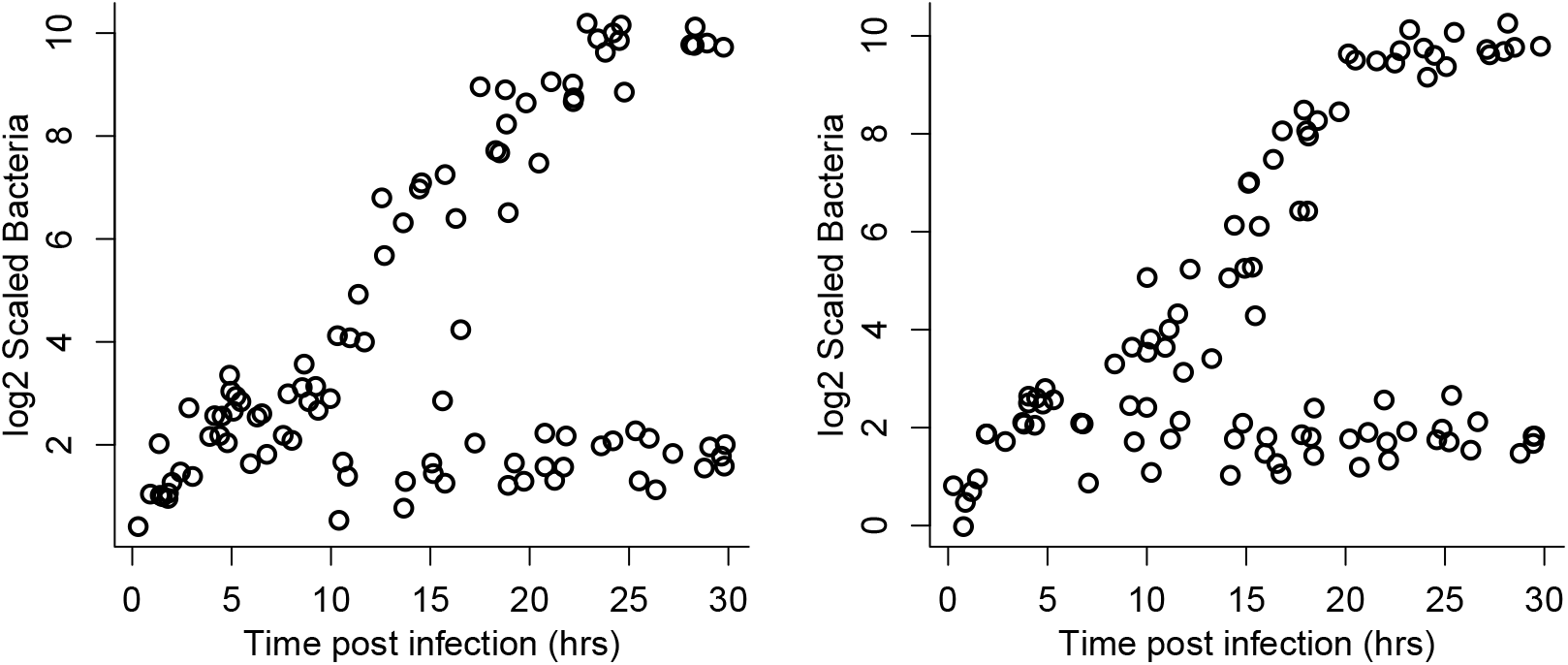
As in Figure 3, for the extended model eqn. (5) with normal and protected bacteria. Parameter values *r* = 0.5; *Q_A_* = 12; *δ_A_* = 0.02; *c* = .1; *K* = 1000; *m* = .25; *δ_R_* = 2; *g* = 0.5; *η* = 0.25; *ν* = 0.5; *L* = 5; *δ_P_* = 0.02 Conversion rate from *N* to *P* was given by the logit function *p*(*A*) = *α*Φ(*A|μ* = *A_P_, σ* = *σ_P_*), where Φ is the cumulative distribution function of a Gaussian distribution with mean *μ* and standard deviation *σ*, with *α* = 0.1, *A_P_* = 5; *σ_P_* = 0.5. Figure made by script `Split_Outcomes_NPAR.R.`

**Figure 5:**
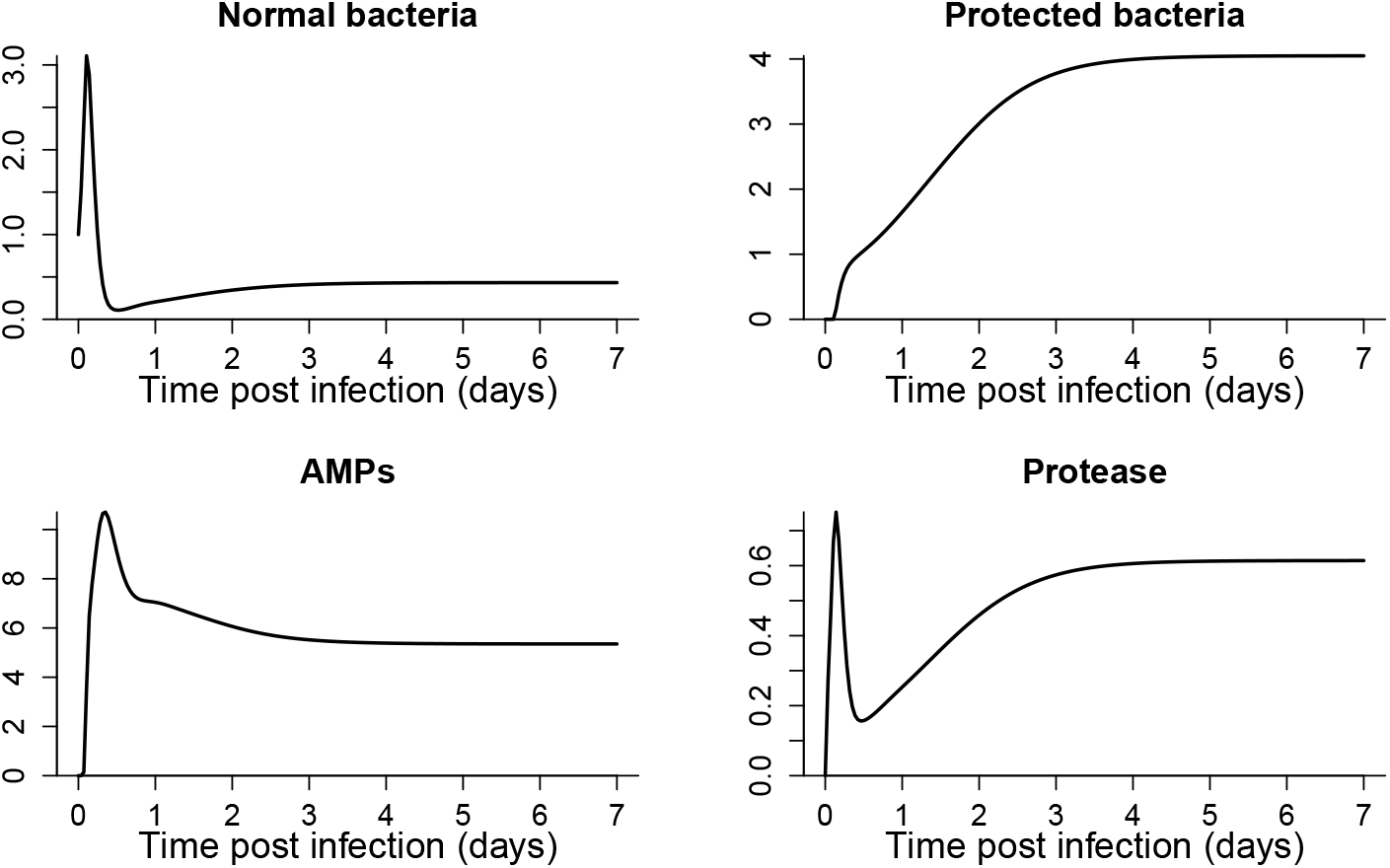
Infection dynamics in the extended model, for a host with fast immune induction so that the outcome is a chronic infection. Parameter values are the same as in fig. 4. Figure made by script `Split_Outcomes_NPAR.R.`

Any observed chronic infection load (those in our *Drosophila* experiments are roughly 10^4^ – 10^5^ per host) can be matched in model (5) through a “protected tissue” scenario where protected bacteria remain near their carrying capacity *L*. But even in this simple model there are multiple ways to achieve any desired equilibrium for *P* as the balance between cell divisions and killing by AMPs.

The outcomes in figs. 4 and 5 are not the only possiblity. In particular, the split into lethal or chronic infections can be transient (Electronic Supplementary Material Fig. S-1). With a larger carrying capacity *L* for the protected pathogens, and sufficiently high conversion rate, a large protected population can become established while the normal, unprotected cells are being driven down by the host immune response. The normal daughters of protected parents can then give the normal population enough of a “boost” that they rebound from near-elimination, and increase to a lethal infection.

## 4 Discussion

The models presented here provide a proof of concept for our general negative feedbacks hypothesis. Hosts mounting a bacteriocidal immune response, and pathogens responding through mechanisms that degrade the bacteriocides or impede their production, is a simple but very general recipe for dynamics where small differences in initial conditions, or small differences between individuals in the values of key parameters, lead to dramatically different outcomes in different individuals. The effects of these differences occur within the first few hours of infection but the ultimate outcome may not be apparent until several days later. When we additionally allow bacteria to enter a protected state at the cost of reduced ability to proliferate, the model can generate outcomes very much like those observed experimentally. The protected state could be a literal safe haven (e.g., a host tissue where they are shielded from immune responses), or a physiological or metabolic state with reduced sensitivity fo the immune response.

Our model for protected pathogens assumed strictly one-way conversion (normal→protected) because that is sufficient to explain chronic infections. Allowing back-conversion would increase the theoretical potential for a suppressed bacteria population to rebound after an initially strong immune response has abated, provoking a second round of immune response. In theory this might lead to cyclical rise and fall of infection, or to a series of infection-suppressiong-rebound events that grow in magnitude and eventually overwhelm the host. However, we are not aware of any empirical evidence for these scenarios in bacterial infections.

Being able to fit previously collected data is not a strong test of a model, especially when information that would constrain model assumptions and parameter values is limited. However, that exercise has produced some new predictions that can be tested experimentally:

1. Chronic infections are dominated by a sub-population of protected pathogens.
2. Protected pathogens are not merely inert and invulnerable - they are doing something that provokes a sustained host immune response. In our models that “something” is that some daughter cells have the normal, unprotected phenotype, but other mechanisms (such as host sensing of metabolic by-products) could have the same effect.

Stronger tests of our hypothesis should involve predicting in advance the outcome of new experiments, using mutant hosts and pathogens with modified kinetic parameters. The models here are built from causal links (e.g., pathogens evoke a host immune response whose strength depends on pathogen abundance) without specifying the underlying “machinery” (e.g., signaling pathways). This is valuable because it means that model predictions are not dependent on those details. However, it does not allow us to test our hypothesis more rigorously by predicting in advance what happens if we monkey with the machinery. To do that, our phenomenological model of the initial activation of host immune responses (a linear ramp from onset to completion) needs to be replaced by a detailed “systems biology” model for the kinetic pathways leading to immune system activation, primarily the Imd signaling pathway. The actual nature of protected pathogens needs to be identified, and state transitions modeled mechanistically. than specifying the outcomes at the level of population parameters (birth, death, and state transition rates). Such models will also help us identify exactly what processes generate the variation among genetically homogeneous hosts, raised in a common environment, that can be amplified into divergent infection outcomes.

## Data accessibility

No original data are presented in this paper. The previously published data used here (in Figure 1), and computer code to replicate all results in the paper, have been uploaded as separate ESM files during submission of the manuscript to make them available to editors and reviewers. Upon publication of the paper, those files will be deposited at figshare or a comparable open archive site.

## Authors’ contribution

All authors contributed to conceiving the study, formulating hypotheses and models, and writing the paper. SPE wrote computer scripts, performed the mathematical analyses, and authored the first draft. All authors gave final approval to submit the paper for publication and agree to be held accountable for the work performed therein.

## Footnotes

Electronic supplementary material is available online at (URL to be inserted at time of publication).

## Electronic Supplementary Material

S. P. Ellner et al., “Host-pathogen Immune Feedbacks Can Explain Widely Divergent Outcomes from Similar Infections”.

**Figure S-1:**
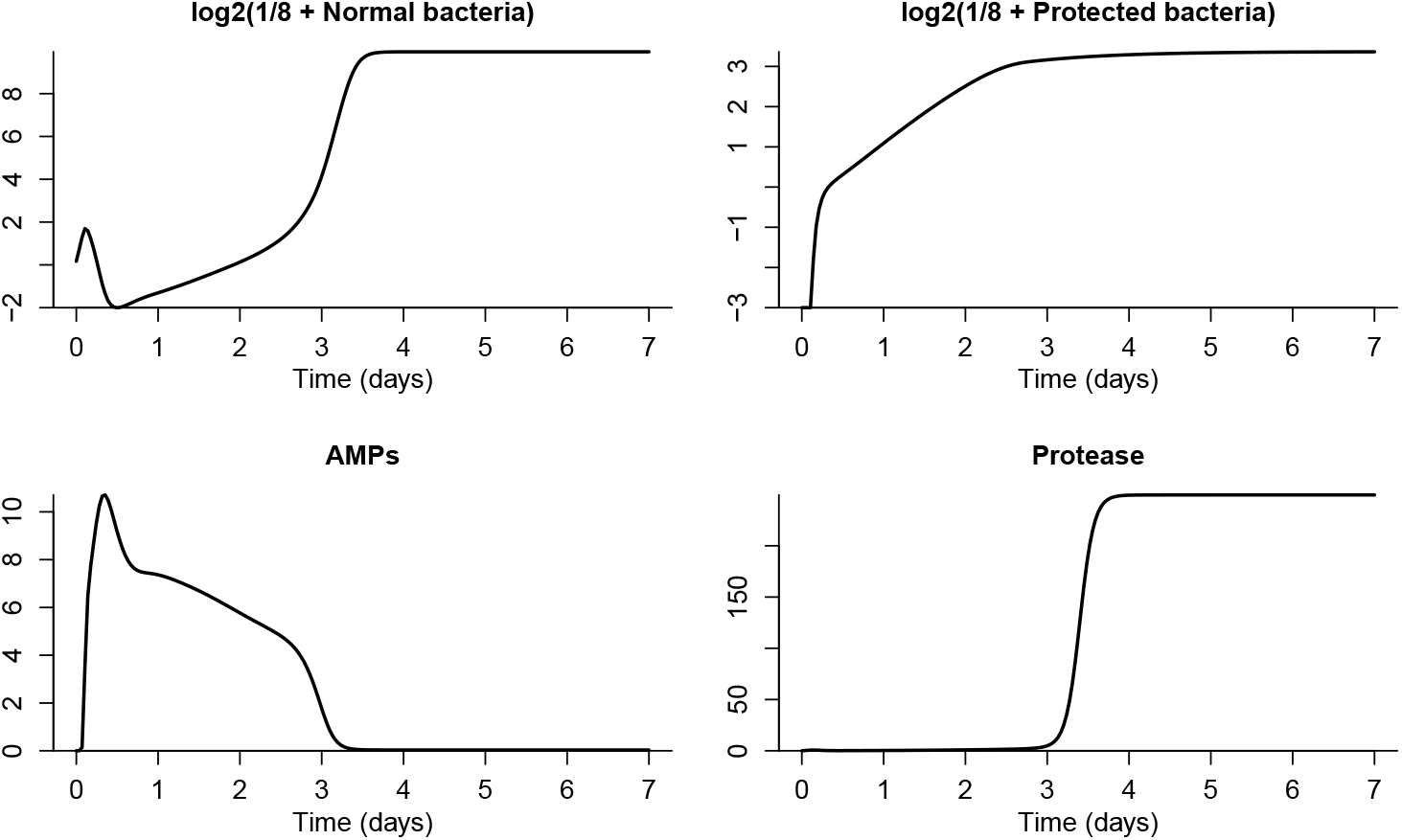
As in fig. 5 except protected bacteria have carrying capacity *L* = 15. Figure made by script `Split_Outcomes_NPAR.R.`

### ESM S.1 Analysis of the basic conceptual model, Eqn. (2)

The analysis uses only standard tools of phase-plane and equilibrium stability analysis. We being with local stability analysis of equilibria, and then demonstrate that periodic and homoclinic orbits cannot occur. We use the notation *ẋ* = *dx/dt*.

There is always an equilibrium is at *B* = *A* = 0. This is the pathogen-free equilibrium, where there are no bacteria, no possibility of bacterial population growth, and no immune response to bacterial infection. Elsewhere on the axes we have 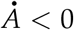 when *B* = 0 and 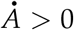 when *A* = 0, so there are no other equilibria on the axes.

Interior equilibria occur at interior intersections of the *B* nullcline *A* = *c*^−1^ (1 − *B/K*) with the *A* nullcline (3). The *A* nullcline begins at *A* = 0 when *B* = 0. Its derivative is positive at *B* = 0 and is easily seen to have a single sign change from positive to negative, being zero at only one point. With increasing *B* the nullcline therefore increases to a unique maximum and then decreases, but it remains positive for all *B* > 0. These properties imply that (as depicted in fig. 2) there is at least one interior equilibrium, and there can be up to three. Because the condition for nullcline intersection is a cubic polynomial, there cannot be more than three.

For stability analysis it is convenient to consider general *f*(*B*) (subject to *f*(0) = 0), and (until further notice) to scale *B* such that *K* = 1. This means that we are scaling *B* relative to the maximum population that could be produced by unchecked proliferation within the host; in the main text *B* is scaled relative to the density that evokes the host immune response at half its maximum possible rate. Many of the calculations are done using MAXIMA in the script file `BAModel.max.`

The Jacobian at (0,0) has +1 as one eigenvalue and the other is negative, implying that the pathogen-free equilibrium is always a saddle. Any interior equilibrium lies on the interior *B* null-cline, where the Jacobian of the model is found to be of the form

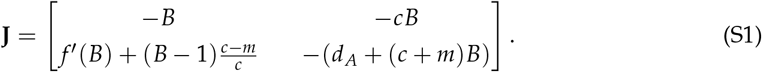

The trace is negative, so stability depends on the determinant,

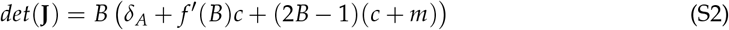

with local stability when *det* (**J**) < 0.

We now show that local stability is determined by the direction of the nullcline crossing at the equilibrium, with stability when the *A* nullcline crosses from below to above the *B* nullcline as *B* increases. This model is simple enough to do the stability calculations explicitly, but we will instead use a general analysis which also covers the extended model in section ESM S.2.

The Jacobian at any internal equilibrium has negative trace (this is shown in scripts `BAmodel.max` and `BAImodel.max` for the present model and the extended model, respectively), so stability depends on the sign of the determinant (stable if positive, a saddle if negative). We can write the model abstractly as

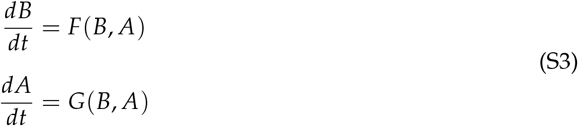

The interior nullclines can be expressed as the graphs of some functions of *B, A* = *Ā*(*B*) and 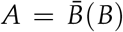, respectively, for the *A* and *B* nullclines. On the nullclines, we have *F* = *G* ≡ 0 and hence (using subscripts to denote partial derivatives)

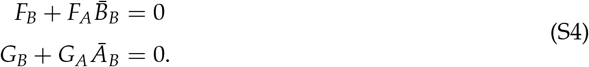

Both of these equalities hold at any interior equilibrium. Solving (S4) for 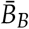 and *Ā_B_*, we have that 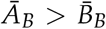 at the equilibrium (i.e., the *A* nullcline crosses the *B* nullcline from below to above as *B* increases) is equivalent to

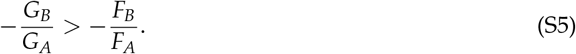

*F_A_* and *G_A_* are both negative in the interior. Multiplying through (S5) by the negative number −*F_A_G_A_* we have *F_A_G_B_* < *F_B_G_A_*, hence *F_B_G_A_* – *F_A_G_B_* > 0. That expression is the determinant of the model’s Jacobian. Hence an “upcrossing” equilibrium is stable, and a “downcrossing” equilibrium is unstable, as illustrated in in fig. 2B. The connection between between nullcline crossing and the Jacobian determinant must be well known; if any reader can give us a citation for it, we would be grateful.

An upcrossing can be either a spiral or a node, depending on the sign of the discriminant 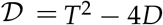 (*T, D* =trace and determinant of the Jacobian). Returning to our conceptual model (2), 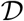 includes a term −4*Bcf*′(*B*) which is ≤ 0 because *f* is by assumption non-decreasing, and no other term involving *f*′ or *f*. Consequently 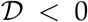 when *f*′(*B*) is sufficiently large, implying complex conjugate eigenvalues, hence the equilibrium is a spiral. On the other hand, suppose the upcrossing occurs when the slope of the *A* nullcline is 0 or smaller. Using (S4), at a tangent intersection of the two nullclines, the determinant of the Jacobian is exactly zero. Therefore, at an upcrossing where the slope of the *A* nullcline is just slightly above the (negative) slope of the *B* nullcline, the determinant will be nearly zero. The trace at an equilibrium equals −*B*(1 + *m* + *c*) – *δ_A_* which is strictly negative (see script `BAmodel.max`), hence the discriminant 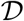 is positive and the equilibrium is a node. This confirms that the transition from three equilibria to one always occurs through a saddle-node collison, as we would expect.

Before returning to our specific model, we note that these general arguments about equilibrium stability and type also apply to the extended model in the following section.

The three-equilibrium case is the one of most interest. Analytic conditions for the three-equilibrium case to occur in our conceptual model involve a cubic polynomical in *B* and so are hard to interpret. However, we can derive an approximate conditionfor the biologically relevant case that *δ_A_* ≪ 1, meaning that little degradation of AMPs occurs on the time scale of interest (hours to days after an infection occcurs). For this calculation we return to the scaling used in the main text, which sets *S_A_* = 1 rather than *K* = 1. Then except when *B* is very small (so that (*c* + *m*)*B* is comparable in magnitude to *δ_A_*) the *A* nullcline (eqn. (3)) is approximately given by

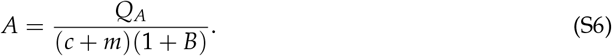

What this approximation misses is that the *A* nullcline actually takes a “nosedive” down to *B* = *A* = 0 as *B* ↓ 0, and if that “nosedive” starts above the *B* nullcline it gives rise to an additional equilibrium with *B* very close to zero.

With a bit of algebra, intersections of the *B* nullcline with the approximate *A* nullcline occur where

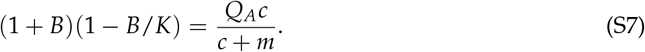

The three-equilibrium case occurs when this quadratic equation has two positive roots sufficiently far from *B* = 0 that eqn. (S6) is accurate, and the “nosedive” equilibrium is then the third equilibrium. The left-hand side of (S7) is a downward-curving parabola with roots at −1 and *K*. There will be two solutions of (S7) with *B* > 0 when the (i) *K* > 1 so that the peak of the parabola occurs when *B* > 0, (ii) the parabola is below the right-hand side at *B* = 0, and (iii) the parabola is above the right-hand side at its peak. These conditions are satisfied iff

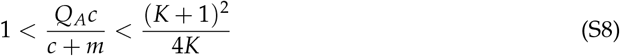

The outer inequality implies *K* > 1, the inner two imply the other conditions. Although (S8) is approximate, we have found that it provides a reliable recipe for finding parameter values at which there are three interior equilibria: pick *c, m* so *c* + *m* is well above *δ_A_*, choose *Q_A_* well above (*m* + *c*)/*c*, and *K* such that 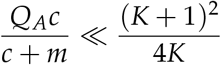.

We now consider global properties of solution trajectories.

First, we find a trapping region in the first quadrant. The axes cannot be crossed from the interior of the first quadrant because {*B* = 0} is invariant, and 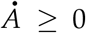 when *A* = 0. Clearly 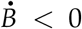 whenever *B* > *K*. When *B* = 0, 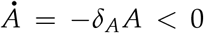 for all *A* > 0, and for *B* ≠ 0 we have 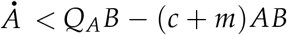 so 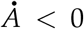 for *A* ≥ *Q_A_*/(*c* + *m*). Therefore the rectangle {0 ≤ *B* ≤ *K*, 0 ≤ *A* ≤ *Q_A_*/(*c* + *m*)} is invariant. Moreover, the derivative bounds imply that eventually *B* ≤ *K* and *A* ≤ *Q_A_* / (*c* + *m*), so that rectangle is eventually entered by any trajectory starting outside it. The portion of the *B* nullcline interior to the first quadrant is a line running from one axis to the other, and any interior equilibrium must lie on this line. By enlarging the rectangle (if necessary) to contain the interior portion of the *B* nullcline, we have an attracting and trapping region that contains all equilibria in the first quadrant.

Second, periodic solutions can be ruled out using the Bendixson-Dulac negative criterion for planar systems. Let *ẋ* denote *dx/dt* for any variable *x*. The criterion says that if there is a smooth function *g*(*B, A*) such that

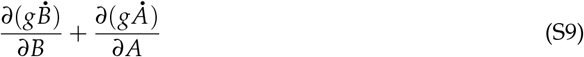

has constant non-zero sign almost everywhere in a simply connected region 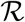 in the plane, then there is no closed orbit contained in 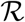. For (2) we consider the region 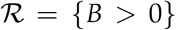 and take *g*(*A, B*) = 1/*B*. With some help from MAXIMA (`script BAmodel.max`) we find that (S9) equals

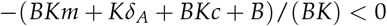

so there are no closed orbits contained entirely in the region *B* > 0. Can there nonetheless be a closed orbit contained in {*B* ≥ 0}? Any such orbit that is not contained in {*B* > 0} must include a point where *B* = 0. But the region *B* = 0 is invariant, hence that orbit must lie entirely in {*B* = 0}, which is impossible. Thus, there cannot be any closed orbits in the region *B* ≥ 0 other than the trivial equilibrium *B* = *A* = 0. Note that the functional form of *f*(*B*) is irrelevant for this result, because *f*(*B*)/*B* is independent of *A* and so it contributes nothing to (S9).

Additionally, there cannot be a homoclinic orbit running from (0,0) to itself. Any such orbit would constitute the stable and unstable manifolds of (0,0). However, the stable manifold of (0,0) is the axis {*B* = 0} on which the dynamics are 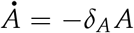, and trajectories on the axis do not approach (0,0) as *t* → −∞.

Finally, we show that when there is only one interior equilibrium, it is globally stable in the interior of the first quadrant. Any solution trajectory eventually enters and stays in the trapping region, so (by the Poincaré-Bendixson Theorem) its *ω*-limit set must be an equilibrium, a periodic orbit, or a finite set of equilibria connected by homoclinic and/or heteroclinic orbits. With only one interior equilibrium, no periodic orbits, and the origin a saddle that can only be approached along an axis, the *ω*-limit set must be either that one equilibrium, or that equilibrium and a homoclinic orbit originating and ending at the equilibrium. However, we have shown above that when there is only one interior equilibrium, it is locally stable, so there cannot be any such homoclinic orbit. Thus, every trajectory originating in the interior of the first quadrant comes arbitrarily close to the unique interior equilibrium, and therefore must converge to it.

### ESM S.2 Analysis of an extended model

As noted in the main text, our model assumes that there is no constitutive defense (AMP production rate is zero when *B* = 0) and the pathogens remove existing AMPs by degradation and sequestration, rather than interfering with their production. To show that our conclusions are not special to that situation, for the moment suppose that there can be a low-level constitutive defense, and that effectors *R* interfere with AMP production, either as an alternative to degradation or in addition to degradation. Specifically, assume that AMP production rate is positive at *B* = 0, and is decreased by a factor *e^−μB^* in the slow-fast reduction of the scaled model where *R* is proportional to *B*. To allow outcomes where the host defense is successful, we assume *μ* = *O*(1) at most. This means that immune suppression does not already stifle AMP production before the pathogen density is high enough to induce an immune response with nontrivial effects on the pathogen. The equation for *A* in the scaled model is then

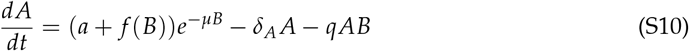

where *f*(*B*) = *Q_A_B*/(1 + *B*), as in the original model, and *q* ≥ 0 represents the combined effects of degradation and sequestration (which in general could be absent). The *A* nullcline is given by the curve

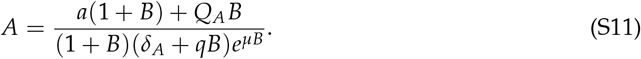

*α* = *a*/*δ_A_* is the constitutive level of defense, i.e. the steady-state value of *A* in the absence of pathogens. The equilibrium *B* = *A* = 0 in our original model thus becomes *B* = 0, *A* = α when constitutive defense is possible. The biologically relevant situation is that *α* is small relative to the values that can occur when pathogen is present. The slope of the nullcline at *B* = 0 is 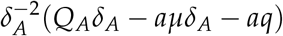 which can be positive or negative at zero.

Elementary calculus shows that the slope of the *A* nullcline is a positive factor times a term that has the sign of the slope at 0, and decreases monotonically in *B*, eventually becoming negative. Thus, when the slope at 0 is positive, the the nullcline has the same qualitative shape as in our original model, increasing at *B* = 0 up to a peak, and then decreasing. When the slope at zero is negative, it remains negative for all *B*.

The Jacobian at the pathogen-free equilibrium (*B* = 0, *A* = *α*) has eigenvalues –*δ_A_*, 1 – *αc* so for *α* small it will be a saddle, as in our original model. Specifically, it will be a saddle whenever the pathogen-free equilibrium lies below the *B* nullcline. Thus, the three qualitative options shown in fig. 2A also hold for the modified model. The Bendixson-Dulac criterion rules out periodic orbits in the region *B* > 0, and periodic or homoclinic orbits that include a point where *B* = 0 are also ruled out, by arguments identical to those for the original model. The general argument in sec. ESM S.1 shows that equilibrium stability depends on the direction of nullcline crossings, exactly as in the original model, and that the type (spiral or node) depends on the steepness of the crossing.

With constitutive defense possible (*a* > 0), all the possible nullcline configurations in the original model (fig. 2A) remain possible, but it is also possible for the pathogen-free equilibrium to lie above the *B* nullcline, as illustrated in fig. S-2A. In that situation, the Jacobian eigenvalues imply that the pathogen-free equilibrium is a stable node. This creates the potential for the bistability scenario shown in which the low-B stable equilibrium has *B* = 0 exactly. If the interior *A* nullcline is entirely above the interior *B* nullcline (not shown), the pathogen-free equilibrium is the only equilibrium and therefore globally stable, for the same reasons as in the original model. However, these cases only occur when the constitutive defense is so strong that a small introduced pathogen population is quickly exterminated.

**Figure S-2:**
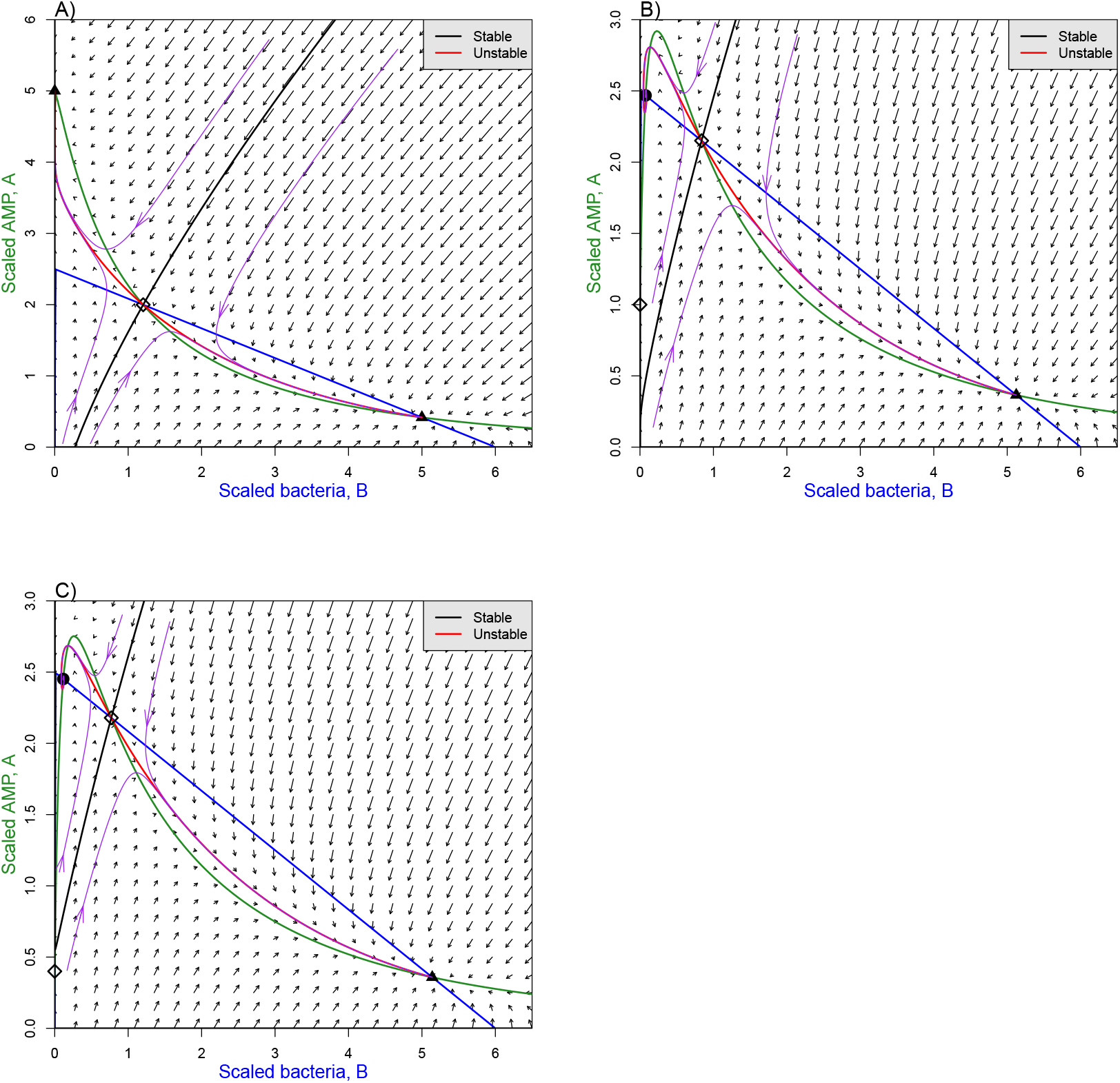
Phase-plane diagrams of the extended conceptual model. **A)** The nullcline configuration producing bistability in the extended model when the constitutive defense equilibrium *B* = 0, *A* = *a/δ_A_* lies above the *B* nullcline. **B),C)** Nullcline and stable manifold configurations that can occur when there is a low-level constitutive defense. Parameter values *Q_A_Q* = 2.4, *δ_A_* = 0.05, *c* = .4, *K* = 6, *q* = .5, *mu* = 0.15 in all panels, and *a* = 0.25,0.05,0.02 in panels A), B), C) respectively. Figures created by script file `BAIModel.R`.

In the more realistic situation of low-level constitutive defense, the extended model offers one more scenario for small between-host variability to produce bimodal outcomes. For parameter values such that the stable manifold of the interior saddle approaches the *A* axis, invasion of the host by a small bacterial population (i.e., initial conditions 0 < *B* ≪ 1, *A* = *α*) lead to chronic infection if the pathogen-free equilibrium is above the stable manifold (fig. S-2B), and lethal infection if the opposite is true (fig. S-2C).

It is reasonable to ask if the extended model model might offer an explanation for chronic infections, so that we do not need to posit protected pathogens, but this cannot occur for biologically reasonable parameter values. That is, it is not possible for the value of *B* at the low-*B* equilibrium to change much. Increasing *a* from zero (as in the original model) to a positive value moves the *A* nullcline up, and decreases the *B* value at the low-*B* equilibrium. Increasing *μ* from zero has the opposite effect, but it is small. The effect can be approximated by using the Implicit Function Theorem to compute the derivative of a nullcline intersection point with respect to perturbation of *mu* away from zero (see script `BAImodel.max`). At *B* = 0 that derivative is zero. The derivative at small *B* is therefore *O*(*B*), hence the change in *B* at the low-B equilibrium is *O*(*μB*). Extremely strong suppression of the host immune response by a small number of bacteria would therefore be required for suppression to have a substantial effect on the location of low-*B* equilibrium.

### ESM S.3 Modeling the Imd signaling pathway

In this section we present the structure and assumptions of our model for the Imd signalling pathway leading to AMP production, and the resulting dynamic equations. We then show that the model can produce a wide range of temporal patterns for the ramp-up of AMP production rate, to justify the simulations of infection dynamics models in the main text where between-host variation in the ramp-up temporal pattern leads to bimodal infection outcomes. Mathematical derivation of the dynamic equations is in Electronic Supplementary Material ESM S.4.

Figure S-3 summarizes our model for the Imd signalling pathway. We assume that the fork of this pathway through PGRP-LE is much less important for immune activation [19], and model the fork through PGRP-LC. Our model is based on the experimentally determined structure of the IMD pathway [13], but as noted below, at one point we simplify the model by assuming that a particular step happens quickly relative to the others. In addition to the initial signaling cascade, our model includes the main known feedbacks whereby up-regulated gene products modify the reaction rates of steps in the cascade, because those can play an important role in shaping the temporal pattern of immune activation (see ESM S.3.1 below).

**Figure S-3:**
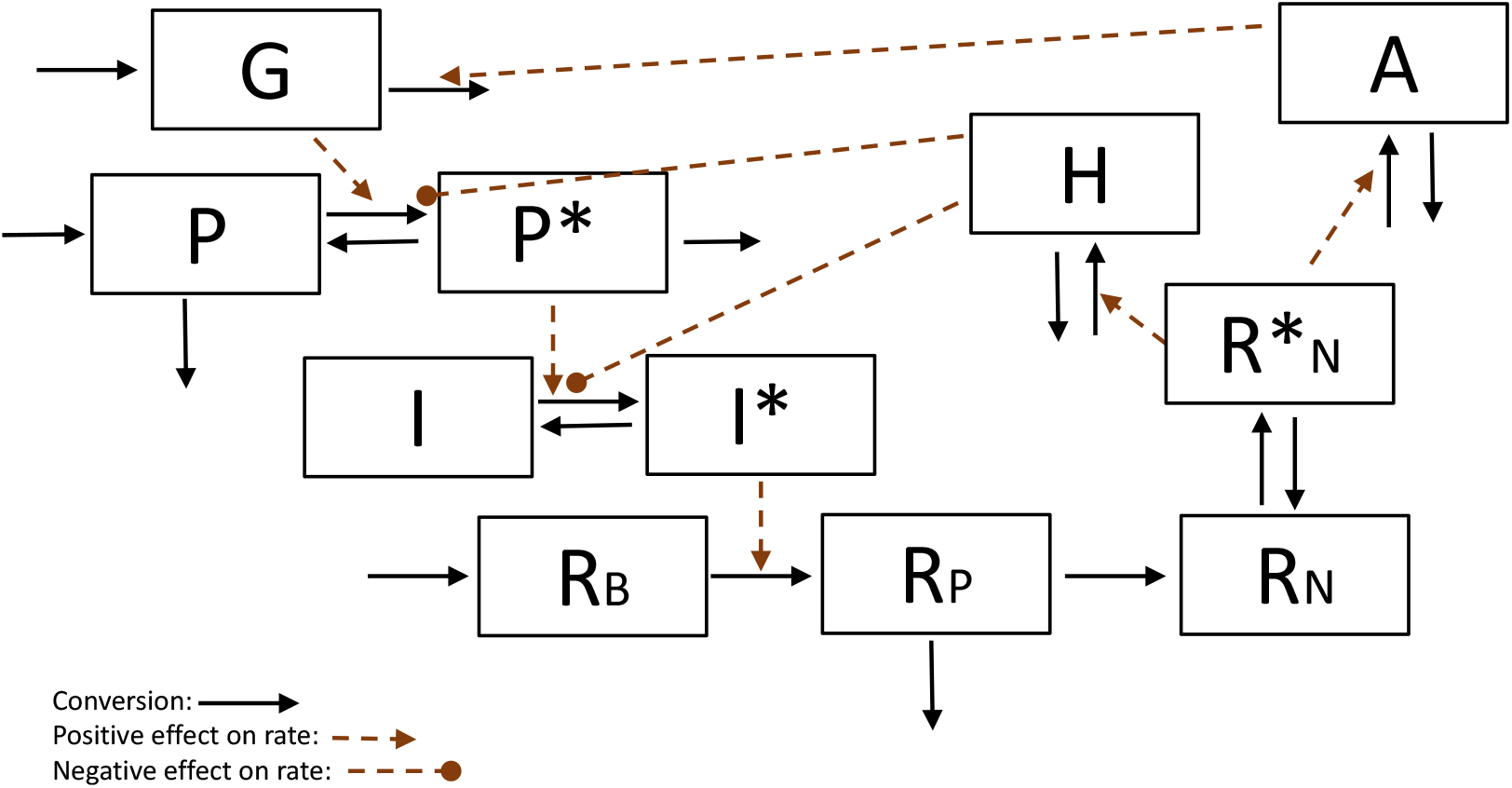
Our simplified model for the fork of the IMd signaling pathway in *D. melanogaster* cells going through PGRP-LC. Solid black arrows indicate flows. Dashed red lines indicate affects of a variable’s concentration on the rate of a reactions, either positive (lines ending in arrowheads) or negative (lines ending in solid circles). The state variables are *G*: peptidoglycans exterior to the cell (moles/l); *P, P*^*^: unbound and bound PGRP-LC (moles); *I, I*^*^: free and recruited IMD (moles); *R_B_, R_P_*: Relish B and P outside the nucleus; • 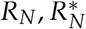: unbound and bound Relish in the nucleus; *H, A*: feedback effector molecules (of various kinds) that act within the cell (*H*) and exterior to the cell *A* (moles). We consider *A* to include AMPs that defend against bacteria.

The steps in our model of the pathway are as follows.

1. Presence of bacteria is indicated by peptidoglycans (PGN) *G* (moles/l), external to the cell. We will eventually couple the Imd pathway to a population of bacteria, in which creation and loss of PGN is explicitly modeled. In this section we treat [*G*] as an exogenous variable affected by feedback effector molecule *A* as described below.
2. Signaling is initiated by PGN *G* binding to PGRP-LC *P* to produce bound PGRP-LC, *P*^*^. We assume that this is a higher-order reaction, as PGN are a polymer and PGRP-LC form polymers on binding to it.
3. Bound *P*^*^ catalyzes the conversion of free Imd *I* to recruited Imd *I*^*^; reversion from recruited to free is assumed to have first-order kinetics.
4. In our model, recruited Imd catalyzes the conversion of Relish_*B*_ to Relish_*P*_. This is a deliberate simplification. In reality, recruited Imd catalyzes the conversion caspase to active caspase (which catalyzes conversion of Relish_*B*_ to Relish_*U*_) and the conversion of kinase to active kinase (which catalyzes the conversion of Relish_*U*_ to Relish_*P*_). Because these steps are not affected by the negative regulators (described below), we can simplify the model without losing any qualitative features by assuming that active caspace and kinase are “readout” variables for recruited Imd, and ignoring the intermediate form Relish_*U*_.
5. Relish_*P*_ can be transported into the nucleus, where it can be unbound (*R_N_*) or bound to promoter sites for the production of AMPs and negative regulators 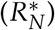. Binding and unbinding are not especially fast relative to other steps in the cascade, so we cannot assume that bound vs. unbound Relish_*P*_ are in steady-state with respect to the total amount in the nucleus. By only tracking the total bound amount, we simplify the reality that nuclear Relish actually binds to several promoter sites for different processes.
6. Similarly, although multiple pathways connect nuclear bound Relish to its feedbacks on the signaling cascade, we aggregate them into two effectors, one acting in the cytoplasm which we call *H*, and the other (called *A* for amydase) which acts outside the nucleus. *H* impedes formation of *P*^*^ and *I*^*^ in the nucleus, and *A* degrades free PGN. We assume that both *A* and *H* are produced even in the absence of bound Relish, but their production rate increases in proportion to the amount of bound Relish.
7. Bound nuclear Relish also up-regulates production of bacteriocidal AMPs. Rather than model this separately, we assume that AMP concentration is proportional to *A*.
8. There is also positive feedback from to production of Relish*B*. If this feedback is directly proportional to the amount of bound nuclear Relish in the nucleus, the model can produce an unlimited spiral of Relish increase. We therefore assume a Michaelis-Menten saturating relationship for this feedback.

The rate equations for each of these reaction steps are derived in Electronic Supplementary Material ESM S.4. Notation for the model is summarized in Table S-1, and the resulting model equations are presented in Table S-2.

#### ESM S.3.1 Temporal patterns of immune activation

We now reach the second aim of this section, which is to explore the range of temporal activation patterns that the model can produce. AMP production rate is proportional to 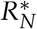, so we ask how 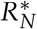 can increase over time from its initial value of zero up to a steady state, when the pathway is activated by *G* going from zero to a positive value.

**Table S-1:** Parameters and state variables for the Imd pathway model and their definitions. Active or bound forms of a molecule are indicated by a star, as in *I*^*^. Coefficient subscripts indicate what they multiply in the model, e.g. *c_I_* and *δ_I_* multiply *I*.

| Parameter | Definition or formula | Units |
| --- | --- | --- |
| $\delta_X$ | First-order degradation rate of molecule $X$ | $\text{time}^{-1}$ |
| $\rho_{X^*}$ | First-order inactivation or unbinding rate of $X^*$ | $\text{time}^{-1}$ |
| $c_X$ | Rate constant for a reaction involving $X$ and possibly others | concentration(s) ×<br>$\text{time}^{-1}$ |
| $Q_X$ | External supply rate, or replenishment rate, of $X$ | moles/time. |
| $\phi_X$ | Coefficient for the strength of a feedback effect of bound Relish | varies; often 1/concentration |
| $[X]$ | Concentration of $X$ (any variable) | moles/l |
| $G$ | Peptidoglycans exterior to the cell | moles/l |
| $P, P^*$ | PGRP-LC — bound and unbound | moles |
| $I, I^*$ | Imd — free and recruited | moles |
| $R_B, R_P$ | Relish — outside the nucleus | moles |
| $R_N, R_N^*$ | Relish – free and bound, in the nucleus | moles |
| $A, H$ | Effector molecules up-regulated by bound nuclear Relish | moles |

**Table S-2:**
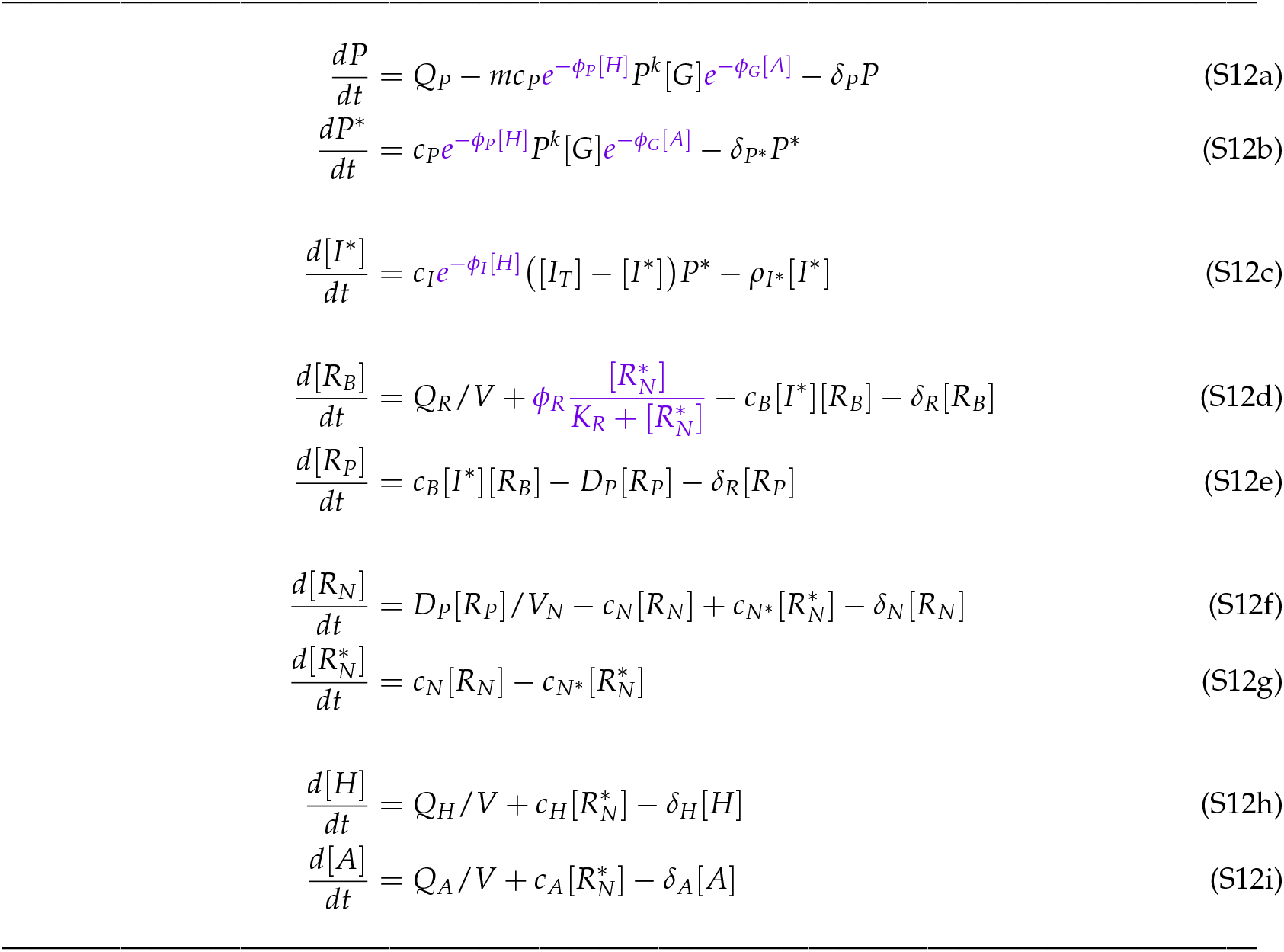
Dynamic equations for the simplified Imd signalling pathway model. The equations assume that all catalyzed reactions are in the first-order phase (i.e., all reactants are at low concentrations) so that saturation, as in the Michaelis-Menten rate equation, can be ignored (Ingalls 2013). Units of state variables are either amounts (moles) in a cell of volume *V*, or a concentration (moles/l) where [*X*] denotes the concentration of *X*. Feedback effects triggered by binding of nuclear Relish are indicated by purple font.

**Table S-3:**
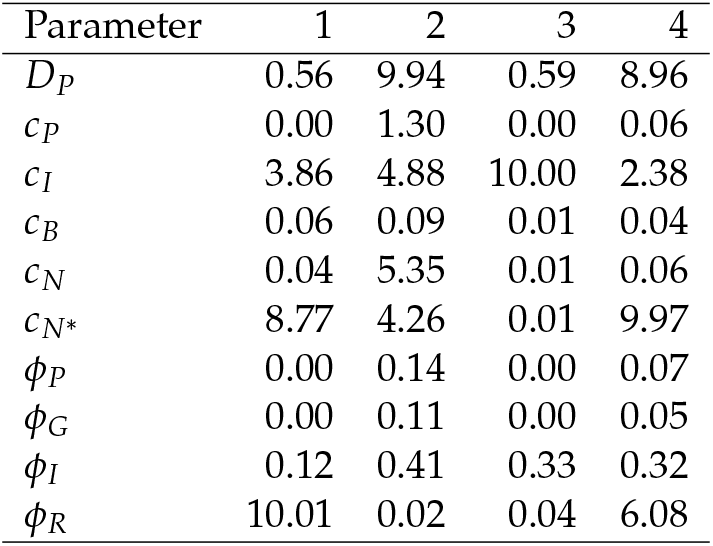
Optimized parameter values corresponding to the four curves in Figure S-4. Columns 1-4 give parameters for the black, red, blue, and purple curves. Other parameters had the same value for all four curves: *V* = 1, *V_N_* = 0.1, *G* = 10, *m* = 2, *k* = 2, *δ*_P_ = .01, *δ*_P*_ = 0.02, *δ_R_* = .01, *δ_N_* = 0.01, *c_H_* = 0.05, *c_A_* = 0.1, *δ_H_* = .05, *δ_A_* = .05, *ρ_I*_* = 0.01, *K_R_* = 2.

The absolute rates of processes in the model are controlled by parameters for which we have no empirical estimates, so we can only ask about relative changes over time. We therefore first rescaled the model so that *P*, [*R_B_*], [*H*] and [*A*] have steady-state value 1 in the absence of pathogen (*G* = 0) and [*I_T_*] = 1. This results in the rescaled model having parameter values

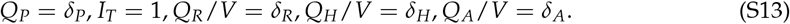

By trial and error, we found a set of plausible “default” parameter values producing a moderately fast ramp-up during a time period centered at time *t* = 4 hours after the pathogen arrives to initiate activation. We then used numerical optimization to find kinetic parameters producing ramp-up patterns that minimized the sum of squared deviations from four “target” patterns of monotone increase from zero to an asymptote (script `Imd_Simplified_RampUp.R`. Values of the parameters *D_P_, c_P_, c_I_, c_B_, c_N_, c_N*_, ϕ_P_, ϕ_G_, ϕ_I_* and *ϕ_R_* were allowed to vary; others were held constant at the default values. The resulting optimal parameter sets included some astronomically large kinetic parameters (up to 10^18^), so we modified the optimization so that the goodness-of-fit was penalized when any parameter exceeded 10 (with time measured in minutes), with the penalty proportional to the square of the excess.

The optimized parameters with that penalty, listed in Table S-3, produced the four curves in Fig. S-4: slow (blue) or fast (red) steady increase starting very soon after the pathogen arrives, or rapid increase following a shorter (purple) or longer (black) delay. Model solutions over a longer time span confirm that all solution curves asymptote to a constant steady-state value. These four do not exhaust the possibilities; they were chosen to illustrate that the model

**Figure S-4:**
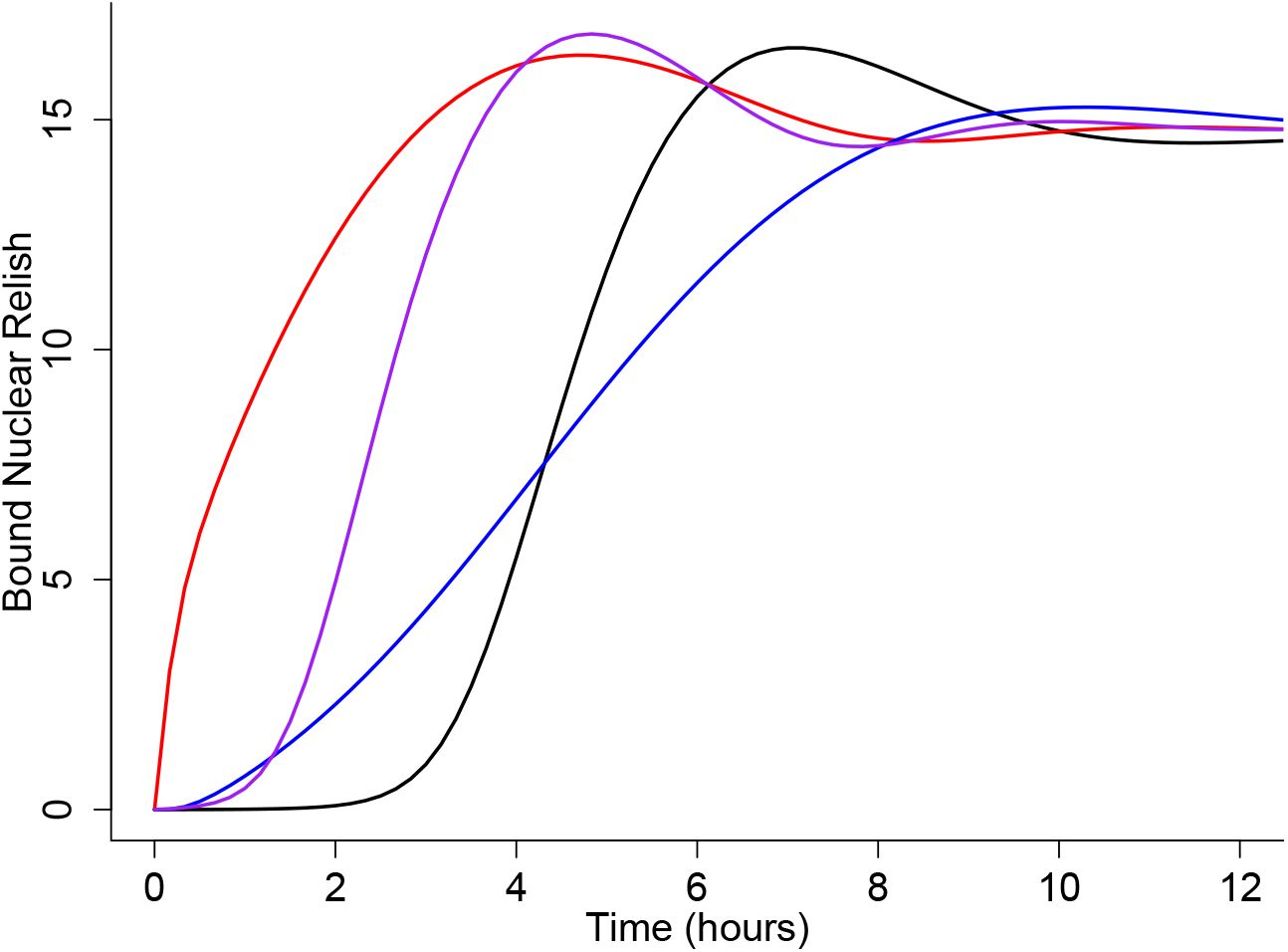
Four possible patterns of immune response activation in the Imd pathway model, Table S-2, rescaled according to eqn. (S13). Parameter values for the curves are given in Table S-3. Figures created by script file `Imd_Simplified_RampUp.R`

### ESM S.4 Derivation of rate equations for Imd pathway model

We consider a signaling cascade initiated by the increase of PGN concentration from 0 to [*G*] > 0. As noted in the main text, we assume that all catalyzed reactions are in the first-order phase that occurs when all reactants are at low concentrations (Ingalls 2013).

Models need to be derived in terms of the amounts of different molecules rather than their concentrations, because amounts flow (e.g., into and out of the nucleus, from bound to unbound states), not concentrations. However, reaction rates per unit volume within in the cell typically depend on concentrations, so many rate equations include concentrations [*X*] = *X/V* (*V* =cell volume), and then the rate per unit volume is scaled back up by cell volume. If the dynamic equation for a extra-nuclear molecule *X* is first-order with respect to [*X*], then the equation can written entirely in terms of [*X*]. That is,

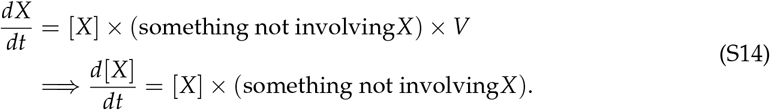

On the other hand, if the reaction is higher-order in [*X*] this generally won’t be true. We do the same for molecules in the nucleus, scaling by nuclear volume *V_N_*.

For transmembrane (PGRP-LC) and within-nucleus molecules, we use amounts rather than concentrations as the units for state variables.

We begin by modeling the initial cascade leading to buildup of Relish in the nucleus. The subsequent feedbacks resulting from bound nuclear Relish will then be added.

The reaction diagrams below often omit degradation processes. To prevent unrealistic buildups, every molecule with a nonzero baseline production rate (in the absence of any stimulus by PGN) is tacitly assumed to have first-order degradation kinetics.

- PGRP-LC, P: 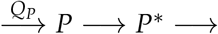 Unbound PGRP-LC is replenished at rate *Q_P_*, and can form a bound complex with PGN. Because PGN is a polymer and a bound complex includes several PGRP-LC molecules, we assume that this reaction rate is higher-order in [*P*] with exponent *k* > 1, and formation of one bound complex eliminates *m* > 1 unbound molecules. A reasonable default assumption is *m* = *k*; this would hold exactly if complex formation involves simultaneous binding of *m* = *k* PGRP-LC molecules, or as an approximation for multistep cooperative binding (Ingalls 2013, sec. 3.3). We assume that there is no reversion from bound to unbound states, but bound and unbound complexes degrade at rates *δ_P_* and *δ_P*_*, respectively. Letting *P*\* denote the number of bound complexes, the kinetic equations are then

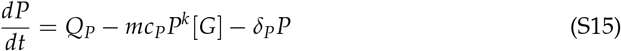

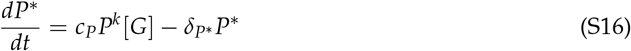
- Imd: *I* ⇌ *I*^*^ Recruitment of free Imd molecules is catalyzed by bound PGRP-LC complexes, and reversion to free Imd has first-order kinetics. Because [*I*] + [*I*\*] remains at some constant level [*I_T_*], we can write an equation for *I*^*^ only:

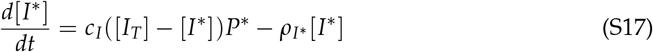
- Extranuclear Relish: 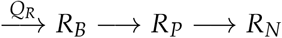 As noted in the main text, *I*^*^ catalyzes formation of activated caspase and kinase, which then catalyze conversion of *R_B_* to *R_U_* and conversion of *R_U_* to *R_P_*. We simplify this step by assuming that the concentrations of activated caspace and kinase are proportional to *I*^*^, and collapsing the conversion of *R_B_* to *R_P_* into a single step. *R_P_* can be transported into the nucleus (*R_P_* in the nucleus is denoted *R_N_*); we assume that this occurs at a rate proportional to its extra-nuclear concentration. This is active transport, rather than diffusion, and we assume that it is irreversible. For simplicity we give the same intrinsic degradation rate *δ_R_* to both forms of Relish.

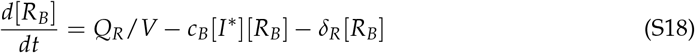

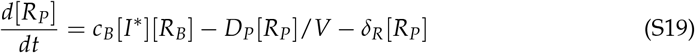
- Relish in the nucleus: 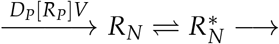 Relish in the nucleus promotes several different processes, so to model in full detail we should consider several different binding sites. However, we simplify this by just classifying Relish in the nucleus as bound or unbound, and have all up-regulated processes respond to the total amount of bound Relish. *R_N_*, 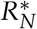 denote unbound and bound, respectively, *R_P_* in the nucleus. Note that Relish diffusing into the nucleus is divided by nuclear volume *V_N_* in the first rate equation because the Relish inflow rate is *D_P_*[*R_P_*].

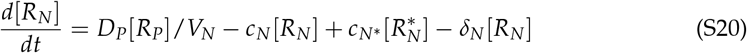

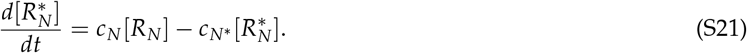
- Effects of bound Relish While multiple pathways connect nuclear bound Relish to its negative feedbacks on the Imd pathway, for simplicity we aggregate them into two effectors, one acting in the cell cytoplasm which we call *H*, and the other (called *A* for amydase) which acts outside the cell. The model tracks their extra-nuclear concentrations:

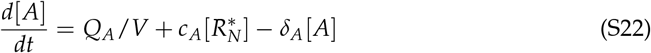

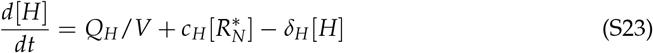 *H* impedes formation of *P*\* and *I*\* in the nucleus, and *A* degrades the external stimulus, free PGN. We assume that both *A* and *H* are produced even in the absence of bound Relish, but their production rate increases in proportion to the amount of bound Relish. Both have first-order degradation kinetics. There is also a positive feedback from bound Relish, an increased production rate of Relish_*B*_. If this feedback is directly proportional to the amount of bound Relish in the nucleus, we can get an unlimited spiral of Relish increase. We therefore assume a Michaelis-Menten saturating relationship.

The final dynamic equations presented in Table S-2 include these feedback effects of *H* and *A*.

